# Single-virion Sequencing of Lamivudine Treated HBV Populations Reveal Population Evolution Dynamics and Demographic History

**DOI:** 10.1101/129023

**Authors:** Yuan O Zhu, Pauline PK Aw, Paola Florez de Sessions, Shuzhen Hong, Lee Xian See, Lewis Z Hong, Andreas Wilm, Chen Hao Li, Stephane Hue, Seng Gee Lim, Niranjan Nagarajan, William F Burkholder, Martin Hibberd

## Abstract

Viral populations are complex, dynamic, and fast evolving. The evolution of groups of closely related viruses in a competitive environment is termed quasispecies. To fully understand the role that quasispecies play in viral evolution, characterizing the trajectories of viral genotypes in an evolving population is the key. In particular, long-range haplotype information for thousands of individual viruses is critical; yet generating this information is non-trivial. Popular deep sequencing methods generate relatively short reads that do not preserve linkage information, while third generation sequencing methods have higher error rates that make detection of low frequency mutations a bioinformatics challenge. Here we applied BAsE-Seq, an Illumina-based single-virion sequencing technology, to eight samples from four chronic hepatitis B (CHB) patients – once before antiviral treatment and once after viral rebound due to resistance. We obtained 248-8,796 single-virion sequences per sample, which allowed us to find evidence for both hard and soft selective sweeps. We were also able to reconstruct population demographic history that was independently verified by clinically collected data. We further verified four of the samples independently on PacBio and Illumina sequencers. Overall, we showed that single-virion sequencing yields insight into viral evolution and population dynamics in an efficient and high throughput manner. We believe that single-virion sequencing is widely applicable to the study of viral evolution in the context of drug resistance, differentiating between soft or hard selective sweeps, and the reconstruction of intra-host viral population demographic history.

## Introduction

Viral intra-host evolution is a critical obstacle in the treatment of infectious diseases. It is the root cause of viral host immune escape and drug resistance, and consequently a major impediment in disease cure and eradication [Hughes and Andersson 2015, Presloid and Novella 2015]. The hepatitis B virus (HBV) is a prime example. HBV is a small, circular DNA virus with a high mutation rate (μ = 10^-5^ to 10^-7^) [Caligiuri et al., 2016]. When coupled with a large viral load (often between ∼10^3^ copies/ml to 10^7^ copies/ml of serum), this can give substantial viral diversity in active infections [Osiowy et al., 2006]. In other words, a sufficiently large viral population can potentially carry, or produce within a short period of time, all possible mutations, thus providing a genetic reservoir for rapid viral response and adaptation [Yim et al., 2006, Margeridon-Thermet et al., 2009, Chen et al., 2009, Tang et al., 2011, Cheng et al., 2012]. Practically, the accumulation of viral mutations is indicative of disease progression and severity [Sterneck et al., 1997, Gunther et al., 1999, Hannoun et al., 2000]. Mutations that quickly become predominant in the population are also indicators for how the viruses might be circumventing host response and treatment that enable fresh approaches for drug development research [Chisari and Ferrari 1995].

Mutations play a vital role in evolution, but the concept of quasispecies is also an important model in viral evolutionary genetics [Eigen 1971, Eigen 1996, Eigen and Schuster 1977, Eigen and Schuster 1978, Eigen et al., 1988, Lauring and Andino 2010]. The term quasispecies initially stemmed from the notion that an evolving population of macromolecules is heterogeneous. Mutation-selection balance in a highly mutable viral population results in populations made up of many similar but non-identical viral sequences, also called mutant clouds. Although the precise definition and exact scope of quasispecies have been debated [Holmes and Moya 2002, Lauring et al., 2013], there is substantial support that some form of quasispecies exists in HBV [Simmonds and Midgley 2002, Osiowy et al., 2006, Domingo et al., 2012]. Having a genetic repository in the form of stable covalently closed circular DNA (cccDNA) in infected hepatocytes ensures the constant presence of ancestral sequence(s), while rapid replication with a high error rate provides the necessary genetic heterogeneity to sustain mutant clouds.

The implications for studying HBV viral evolution are as follows. First, consensus sequence changes occur relatively slowly. For HBV, the mean number of nucleotide substitutions is only estimated at between 1.5 × 10^-5^ to 7.9 × 10^-5^ nucleotide substitutions per site per year [Osiowy et al., 2006]. The study of consensus sequences alone may not reveal underlying quasispecies dynamics, which may be much more rapid as the population constantly explores possible genotypes [Domingo et al., 2012]. Second, these hidden quasispecies dynamics may be important in understanding the key indicators of viral fitness. Human host immune response, host genetics, treatment regimes, and finally the viral genotype itself likely interact in a complex fashion that exerts multiple, possibly contradictory selective forces on the virus that ultimately culminates in clinical outcome. Identifying the relevant subpopulation of viruses that are reacting to selective pressures of interest, whether it is nucleoside analogues, interferon treatments, or a change in host immune response can reveal important viral indicators for disease progression.

In order to leverage the recent advancements in next generation sequencing (NGS) technology, we explored single-virion sequencing as an option for characterizing quasispecies diversity in active infections. Deep population sequencing is routinely used to identify polymorphisms, including extremely rare alleles [Widasari et al., 2014, Yamani et al., 2015, Yan et al., 2015, Li et al., 2015, Ode et al., 2015, Chen et al., 2016]. However, without linkage information, it remains difficult to describe quasispecies based on allele frequencies alone. A large number of complete genomes from a single viral population must be sequenced to be confident of full quasispecies diversity.

Traditionally, such studies require viruses to be individually cloned and sequenced – a tedious, non-scalable process requiring a large amount of work and precious source material [Osiowy et al., 2006, Lim et al., 2007]. Quite often, quasispecies are simply left out in the study of viral evolution. However, the complexity and importance of quasispecies has never been clearer [Andino and Domingo 2015], and there are two recent next NGS technologies that can be applied to single-virion sequencing in a high-throughput manner, promising up to thousands of viral sequences from every chronic hepatitis B (CHB) patient sample. BAsE-Seq is an Illumina-based method that makes use of random 20mer barcodes to tag every single viral genome with a unique sequence. The barcoded genomes are then amplified as a single amplicon for library construction [Hong et al., 2014]. Reads from BAsE-Seq libraries can be reassembled into individual viral genomes *in silico* post sequencing, effectively constructing thousands of viral genomes with full haplotype information. An alternative approach uses single molecule real time sequencing technology (SMRT) on the Pacific Biosciences platform (PacBio) to produce long reads for individual molecules (up to 60kb). While single pass sequencing error rates are high, the relatively small 3kb HBV genomes can be read multiple times by the same polymerase, sharply lowering error rates and yielding highly accurate genome sequences, with the additional benefit of not requiring a reference genome [Dilernia et al., 2015].

We aimed to apply these single-virion sequencing methods in a manner tailored to characterizing viral population diversity, quasispecies structure, and population evolution. More specifically, we aimed to discover additional information on viral evolutionary dynamics not visible to regular deep sequencing that is now detectable with single-virion sequencing. We picked a relatively well-understood model – that of HBV resistance to the antiviral drug Lamivudine, where the most common resistance alleles are well characterized [Pallier et al., 2006], and obtained two serum samples from each of four CHB patients who were treated with and subsequently developed resistance to Lamivudine. We searched for resistance mutations in each of the patients and tried to reveal additional quasispecies dynamics using single-virion sequencing. We found that single-virion sequencing reveals vital information about viral population heterogeneity and fluctuations in population composition during viral evolution.

## Material and methods

### Sample Identification and Collection in the Clinic

Viral DNA was extracted from 200 μl of patient serum using the Qiagen Blood mini kit. The extracted HBV genome was PCR amplified using Dynazyme DNA polymerase and the primers [Fwd (5’-G[T/C]GTAGACTCGTGGTGGACTTCTCTC-3’) Rev (5’-TGACA[T/A/G/C]ACTTTCCAATCA AT-3’)]. The amplified 650bp fragment was purified by gel electrophoresis and extraction and directly sequenced on an ABI 3730XL DNA Analyzers to identify resistance alleles (SI Table 1). Plasmids with clones HBV sequences (referred to as Clone-1 and Clone-2 in the text) were constructed and processed as previously detailed in [Hong et al., 2014].

### Illumina Library Construction and Sequencing

For a detailed protocol for sample library preparation, refer to [Aw et al., 2014]. Briefly, 10^6^ HBV viral genomes were PCR amplified using custom primers that cover all but the first 40bp. 2-3 μg of PCR product for each viral DNA sample was sheared to achieve a peak size range of 100–300 bp. Library preparation was performed using the Qiagen GeneRead DNA Library I Kit according to manufacturer instructions. After end-repair, A-tailing, and adapter ligation, ligated products in the 200 - 400 bp range were gel-extracted, and subjected to 14 PCR cycles to incorporate multiplexing indices. The final product was quantified and run on a Illumina HiSeq instrument. Resulting Illumina 2×101 bp reads were trimmed by base quality and mapped to the concatenated HBV pan-genome consisting of all 8 major genotypes A-H (SI Figure 1, SI Table 2). All concordantly mapped read pairs were duplicate-marked, realigned, and recalibrated with GATK 2.7 [McKenna et al., 2010]. SNVs present in the pool were called based on comparison with the best match genotype sequence using LoFreq 2.1.2 with primer regions masked [Wilm et al., 2012].

### Barcode-directed Assembly for Extra-long Sequences (BAsE-Seq)

Library preparation was carried out according to the protocol as described in [Hong et al., 2014]. Briefly, a total of 10^6^ HBV genomes were subjected to a 2-cycle PCR that assigned unique barcodes to each strand of the HBV genome. Two rounds of PCR were carried out to amplify the product, using HBV specific primers (5′-GCTCTTCTTTTTCACCTCTGCCTAATCA-3′ and 5′-GCTCTTCAAAAAGTTGCATGGTGCTGG-3′), taking care to stay within the exponential amplification regime during each round of PCR to minimize the generation of chimeric PCR products. Samples were exonuclease-digested to generate a pool of nested deletions, end-repaired, and circularized. Circular products were fragmented and tagged with the Illumina adaptors followed by 14 cycles of PCR to incorporate primers for sequencing. The resulting 2×101bp reads were processed with a custom pipeline. In the pipeline, FastQ reads were first trimmed for adaptor sequences and base quality.

Reads were then BWA-MEM mapped to the closest genotype reference [Li and Durbin 2009, Li and Durbin 2010]. Aligned reads were duplicate-marked, realigned, recalibrated, and SNVs were called with LoFreq for incorporation into the viral sequence (SI Figure 4,5,8). Viral sequences that passed all quality filters were written into original SAM files as long reads that map to position 1 of the chosen reference for ease of analysis and comparison across platforms. LoFreq, after masking primer regions, was used to call segregating sites within the population. PHYLIP Neighbor Joining trees were constructed [Price et al., 2009, Price et al., 2010] and plotted on iTOL [Letunic and Bork 2007, Letunic and Bork 2011]. For further details about the pipeline refer to [SI].

### PacBio Library Construction and Analysis

10^6^ HBV viral genomes were PCR amplified using the same custom primers as mentioned above under Illumina Sequencing. 2-3 μg of PCR product was used for PacBio library construction following the 2kb Template Preparation and Sequencing protocol. Library products were quantified on Agilent 2100 Bioanalyzer, and run on a PacBio instrument with V6 chemistry. PacBio raw reads were first processed with the SMRT Portal analysis programs. Circular consensus sequences (CCS) from each library were called with a stringent cutoff of at least 10x subreads within a polymerase read and a minimum subread length of 2500bp using the RS_ReadsOfInsert application (SI Figure 2a,b). CCSs were multiple-sequence aligned against all 8 genotypes with MUSCLE [Edgar 2004], followed by neighbor joining tree construction with PHYLIP and tree plotting on iTOL. Bases within the CCS reads with quality scores <75 were masked as Ns to filter out false positives (SI Figure 2c), and the resulting (nearly) full-length viral sequences were BWA-SW mapped as extremely long reads to the concatenated HBV pan-genome consisting of all 8 major genotypes A-H (SI Figure 3). (Although a reference panel is not necessary for PacBio long reads, it was included in the analysis here for direct comparison between outputs from the platforms.) Segregating sites within the viral populations were called with LoFreq with primer regions masked. For a detailed protocol regarding PacBio read processing and error filters refer to [SI].

### Reconstruction of Demographic History by BEAST

A Bayesian Markov Chain Monte Carlo (MCMC) approach was implemented using BEAST v1.8.4 [Drummond and Rambaut 2007] on all sets of 4 patient samples in order to estimate demographic and evolutionary parameters, using the Bayesian skyline plot as a coalescent prior. Unique single-virion sequences constructed from BAsE-Seq libraries often carried missing information due to uneven coverage. Because an excess of ‘N’s can overwhelm the true signal, only the top 100 sequences with the highest overall coverage were used for BEAST analysis. A final fragment of 3134 coding bases was used for demographic history reconstruction. Samples prior to Lamivudine treatment were defined as sequences collected on day 0 and samples post drug resistance annotated as sequences collected *n* days after. We employed the GTR+Γ_4_ unlinked codon model of nucleotide substitution and a strict molecular clock. The MCMC chain length was set to 1E9 to 2E9 generations, depending on the patient sample in question, with sampling of every 1E4^th^. Convergence of the estimates was considered satisfactory when the effective sample size (ESS), calculated in Tracer v1.6, was >200 for all parameters. The first 10% of the estimates was discarded as burn-in. Where necessary, multiple runs were merged using LogCombiner as part of the BEAST package. Run results were analyzed and skyline plots, showing changes in effective population time over time, generated with Tracer v1.6 [Rambaut et al., 2014].

## Results and Discussion

### Deciding On An Appropriate Platform

The three platforms - Illumina (pooled deep sequencing), BAsE-Seq, and PacBio - were tested on two HBV clones with known sequences [Hong et al., 2014] for pipeline construction and optimization. Pipelines tailored to each platform were then applied to viral populations from two patients (P1 and P2) to gauge single-virion sequencing performance and throughput on clinical samples [SI]. BAsE-Seq was picked as the most appropriate platform for these particular samples due to three reasons – the availability of high quality reference genomes, the low incidence of indels in viral sequences, and higher base quality in Illumina reads as compared to PacBio reads. Based on sequencing runs of known clones, base error rates of BAsE-Seq and PacBio libraries were between 0.02-0.3 and 0.2-1.3 per kb of single-virion sequence respectively. However, the error rates for small indels were <0.02/kb and 2.9-3.4/kb respectively. The small indel error rate in PacBio reads, likely due to polymerase slippage, was a cause for concern. While PacBio error rates could be reduced through careful selection of PCR polymerase [SI Fig 2a-b] and CCS quality filters [SI Fig 2c], resulting reads still faced multiple sequence alignment issues that gave false positive single nucleotide variants (SNVs) at the time of analysis [SI Fig 3]. We bypassed this issue by making use of the available reference sequences to map PacBio reads, with the additional bonus of ease of comparison across platforms due to similar analysis pipelines (SI 6-7, 9-15). However, this also resulted in a protocol that neither required nor fully utilized the advantages of a reference independent sequencing technology. As such, BAsE-Seq with its lower error rates overall became the appropriate choice. A more detailed explanation of all work conducted in this comparison exercise is available in Supplementary Materials.

### Classic Resistance Mutations Observed

Serum samples were taken from each patient twice for viral DNA library construction - once before they were treated with Lamivudine (labeled as P1.1, P2.1, P7.1, P11.1) and once after viral loads rebounded to detectable levels (labeled as P1.2, P2.2, P7.2, P11.2) (Table 1).

**Table 1.** Patient sample nomenclature and viral copy number. ^†^-total number of single-virion sequences obtained from a BAsE-Seq library. *-total number of single-virion sequences obtained from a PacBio library. Nucleotide diversity Π, the arithmetic mean between all pairwise differences between viral sequences within each viral population, were calculated from BAsE-Seq single-virion sequences for the entire amplified genomic sequence of 3,175 bases (3215 minus the 40 bases that were not amplified).

| Sample ID | Patient | Date | Viral Copies/ul | Single-virion Sequences | Nucleotide Diversity $\Pi$ |
| --- | --- | --- | --- | --- | --- |
| 1.1 | 1 | 15<sup>th</sup> Nov 94 | 378,500 | 1717<sup>†</sup>, 1635* | 0.0012/base |
| 1.2 | 1 | 3<sup>rd</sup> Jul 99 | 52,270 | 3331<sup>†</sup>, 2514* | 0.0024/base |
| 2.1 | 2 | 22<sup>nd</sup> May 95 | 239,750 | 391<sup>†</sup>, 1330* | 0.0032/base |
| 2.2 | 2 | 29<sup>th</sup> Aug 97 | 44,687 | 3747<sup>†</sup>, 2504* | 0.0013/base |
| 7.1 | 7 | 19<sup>th</sup> May 95 | 69,180 | 2647<sup>†</sup> | 0.0022/base |
| 7.2 | 7 | 11<sup>th</sup> Apr 00 | 48,083 | 789<sup>†</sup> | 0.0016/base |
| 11.1 | 11 | 19<sup>th</sup> Jun 95 | 208,275 | 2754<sup>†</sup> | 0.0211/base |
| 11.2 | 11 | 3<sup>rd</sup> Nov 98 | 466,200 | 248<sup>†</sup> | 0.0052/base |

The four libraries from patients P1 and P2 were sequenced on Illumina, BAsE-Seq, and PacBio platforms. The remaining four libraries from patients P7 and P11 were sequenced and analyzed only by BAsE-Seq due to limited patient serum availability. Consensus viral genomes were constructed from each platform independently (Figure 1).

**Figure 1.**
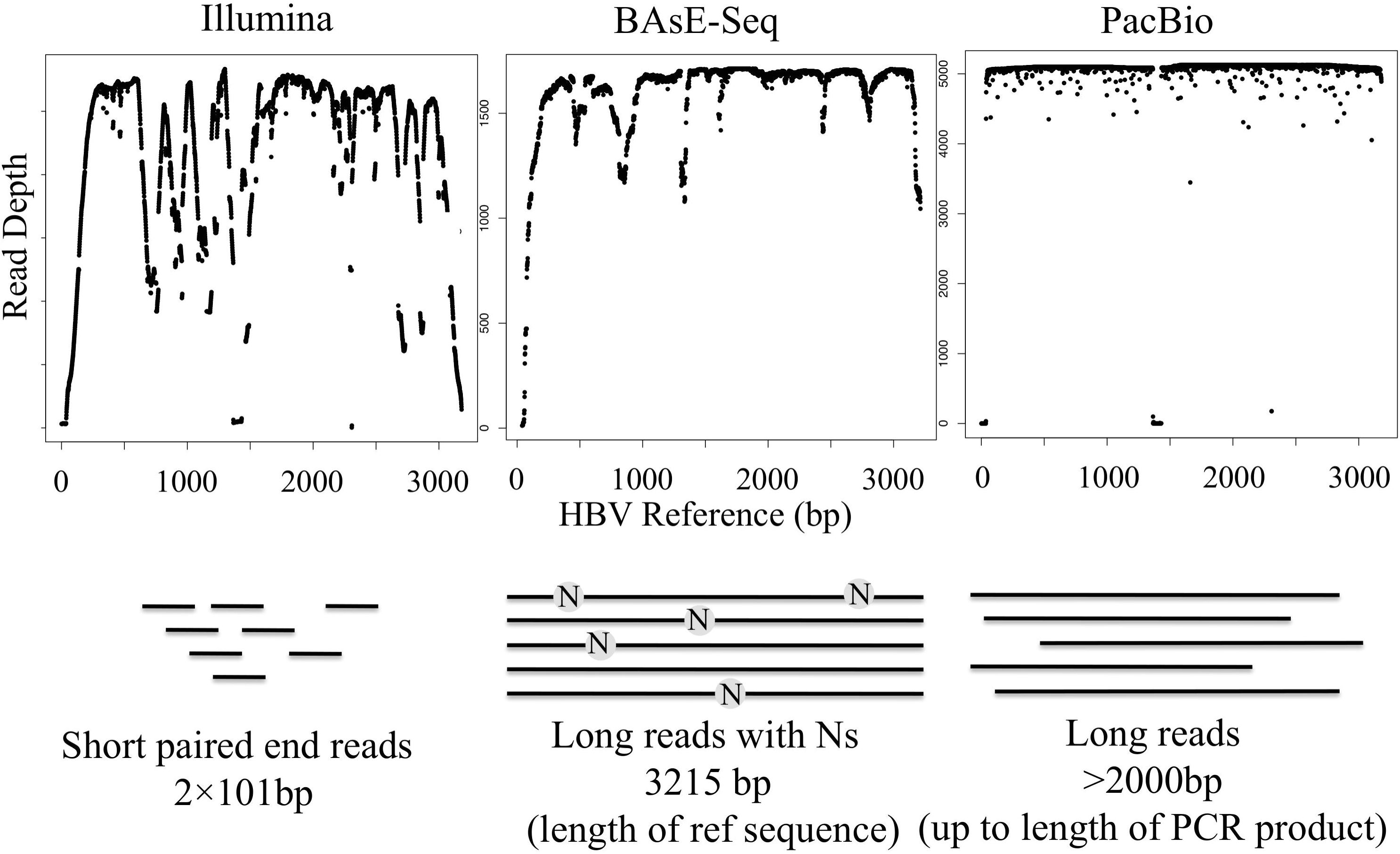
Graphical representation of sequence reads obtained from BAsE-Seq, PacBio, and Illumina platforms. From left to right: Illumina short reads give uneven genomic coverage. BAsE-Seq reconstitutes single-virion sequences by mapping barcoded short reads to a reference, thus matching exact reference length but may have missing information spread throughout depending on local coverage as shown by dips in the coverage plot. PacBio circular consensus reads (CCSs) can vary in length, but will not require a reference for construction, although they are mapped here for comparison. A library of 3kb amplicons will cover the entire genome evenly as shown. Dip in coverage around 1.5kb is due to a deletion in the sample.

Viral genotype composition in each sample was estimated from the percentage of reads mapping to each genotype reference in the pan-genome panel. Three out of four patients carried Genotype B viruses. The only exception was P11, who carried a mixed Genotype B and Genotype C infection prior to drug treatment, but only Genotype C viruses post Lamivudine resistance (Figure 2). Illumina short reads tend to mis-map in regions where sequence divergence is ∼3% between the references used, an issue absent in BAsE-Seq and PacBio long reads [SI Fig 4]. Therefore, genotype identification in Illumina libraries must take into account evenness of coverage across the references, or number of mismatches in mapped reads, in addition to absolute percentage of reads mapped.

**Figure 2.**
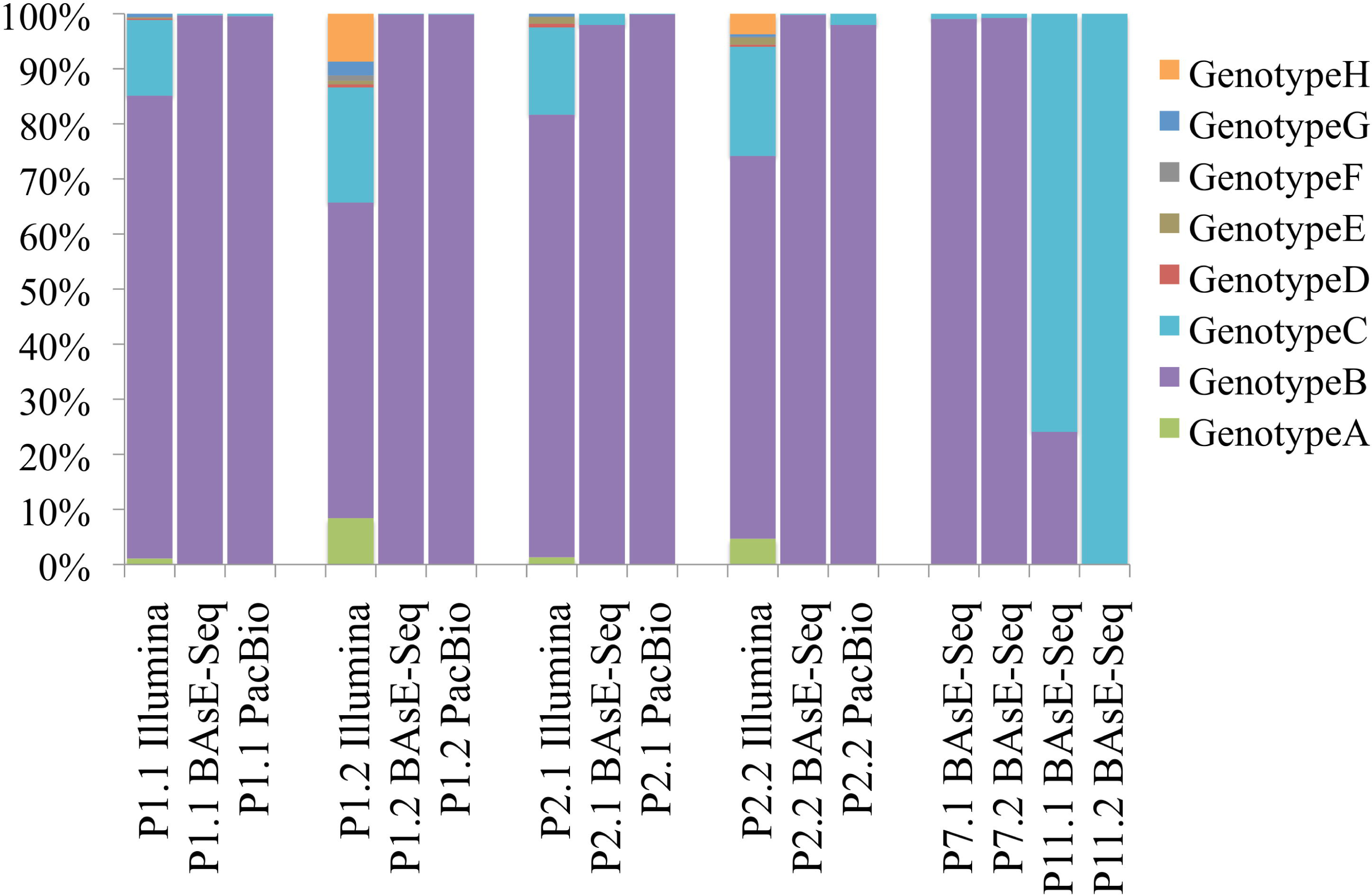
Genotype composition (Y-axis) of samples sequenced (X-axis) as reported by BWA-SW coverage across 8 reference genotype sequences. Samples are organized from left to right in order of P1.1, P1.2, P2.1, P2.2, P7.1, P7.2, P11.1, and P11.2, where P1.1 and P1.2 are two time points from the same patient P1, before and after drug resistance respectively. Wherever data from multiple technologies are available, order follows Illumina, BAsE-Seq, and lastly PacBio.

Lamivudine resistance is achieved through mutations in the RT domain of the polymerase gene in HBV [Pallier et al., 2006]. Two resistance phenotypes made up of three amino acid changes, M204I and L180M+M204V, are the most commonly observed. They confer similarly high resistance and only require one to two nucleotide changes. Both of these resistant genotypes were found, and together explained resistance in all four patients (Table 2). The discrepancy in allele frequencies between the platforms Illumina and BAsE-Seq/PacBio for P1 may have been due to sampling error of a low viral load sample, or mapping errors with Illumina sequencing. Unfortunately we had no remaining patient sample to allow further validation.

**Table 2.** The frequencies of detected resistance alleles in each of the 4 patients after drug resistance. None of these mutations were observed (below detection limit) in the populations prior to development of resistance.

| # | Mutation | Illumina | BAsE-Seq | PacBio |
| --- | --- | --- | --- | --- |
| P1 | M204V | 0.174 | 0.711 | 0.567 |
|  | L180M | 0.165 | 0.752 | 0.611 |
|  | M204I | 0.803 | 0.251 | 0.358 |
| P2 | M204I | 0.981 | 0.882 | 0.990 |
| P7 | M204V |  | 0.870 |  |
|  | L180M |  | 0.946 |  |
| P11 | M204I |  | 0.948 |  |
|  | L180M |  | 0.679 |  |

While P2 (M204I) and P7 (M204V+ L180M) carried single resistance phenotypes, P1 carried both M204I and M204V+ L180M. The genotypes are not mutually exclusive and all three of the point mutations were found in the same patient at significant frequencies (Figure 3,4). P11 was nearly fixed for M204I, but also carried L180M at a high frequency.

**Figure 3.**
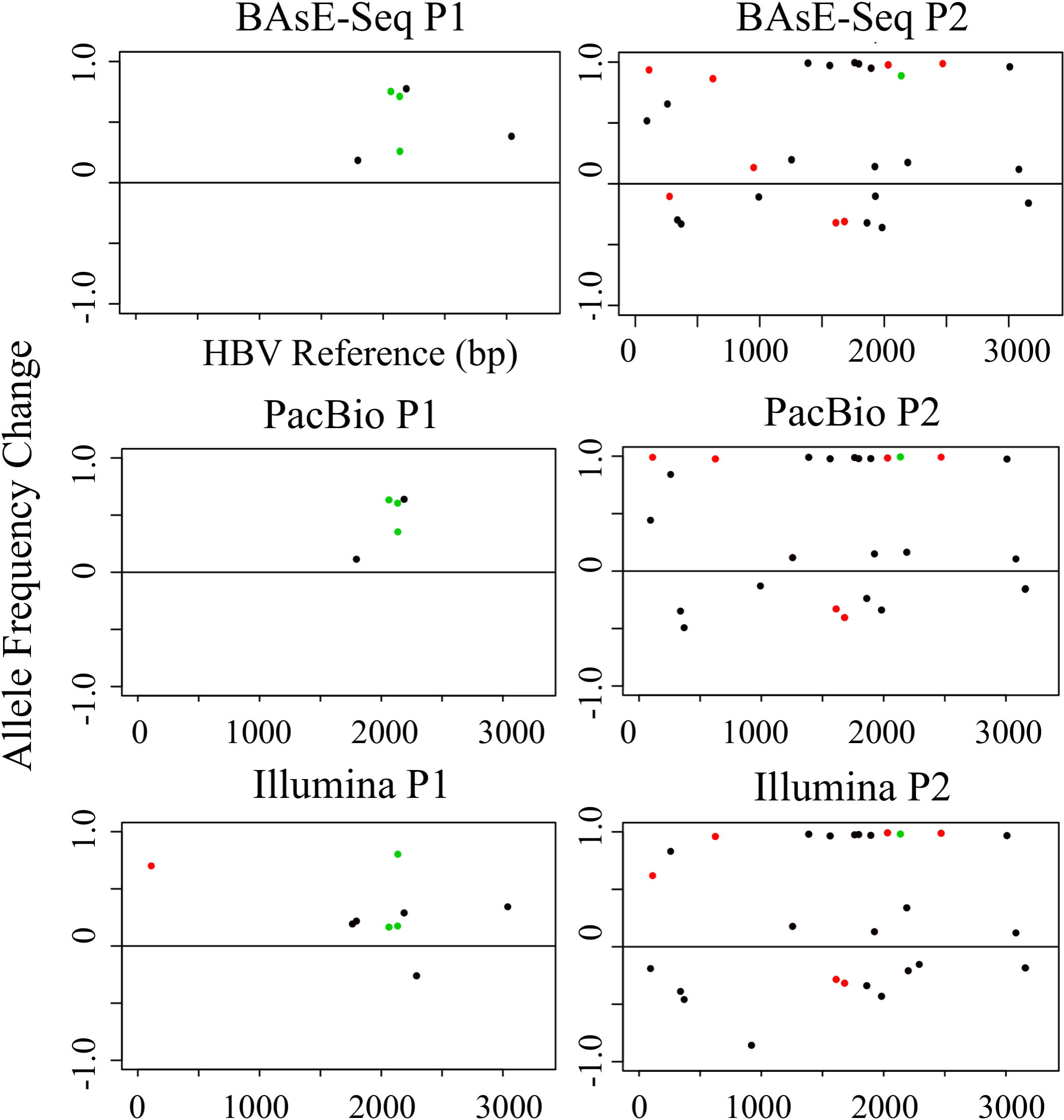
Change in allele frequencies of variable sites in P1 and P2 after drug treatment (detection limit >0.01). Left panels shows the delta change in allele frequencies of segregating sites (Y-axis) across the genome (X-axis) in patient 1 on all 3 sequencing platforms. Right panels show the corresponding data for patient 2. All non-synonymous changes were colored in black, synonymous changes in red, and known resistance alleles in green.

**Figure 4.**
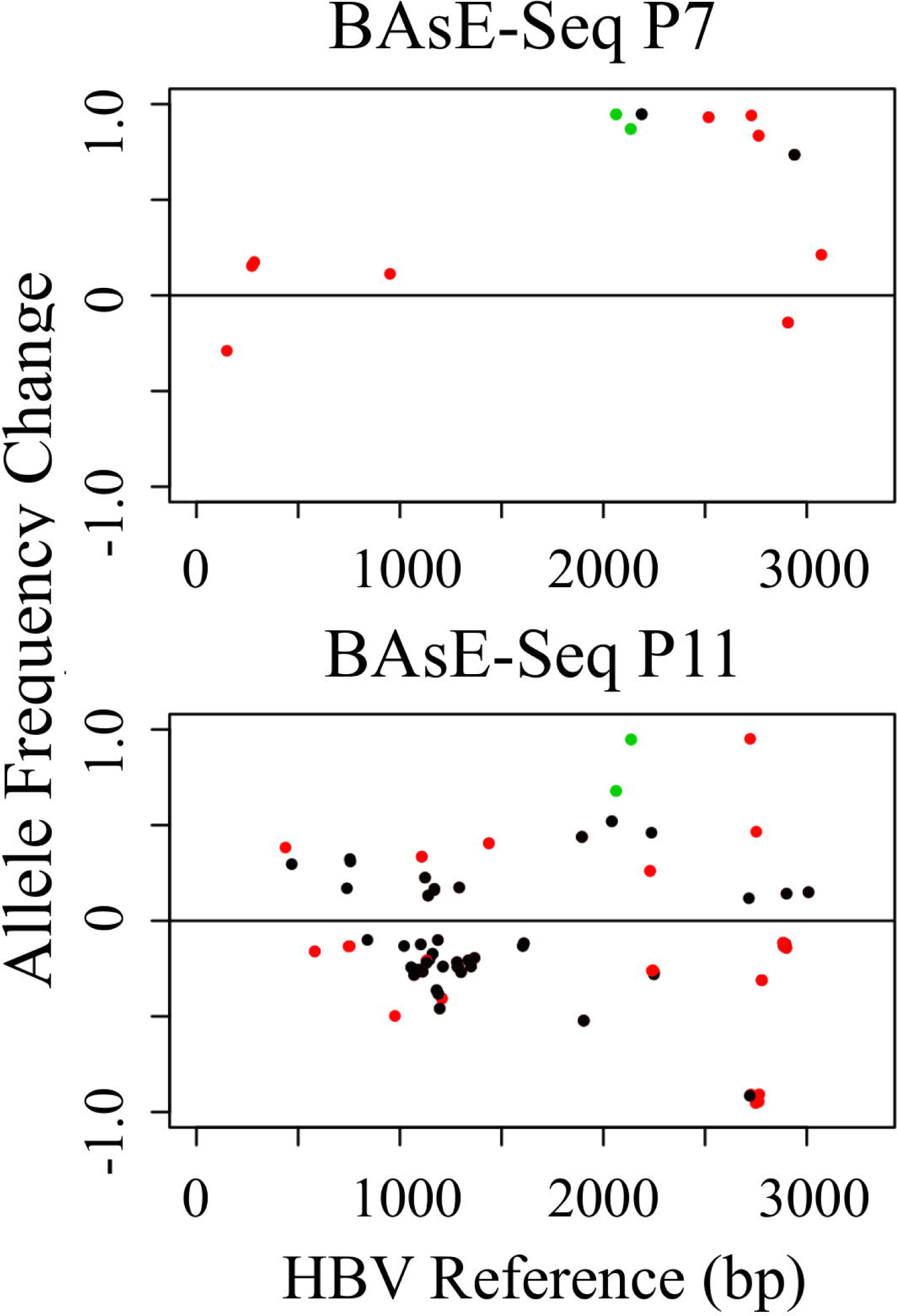
Change in allele frequencies of variable sites in P7 and P11 after drug treatment (detection limit >0.01). Panels show the delta change in allele frequencies of segregating sites (Y-axis) across the genome (X-axis) from BAsE-Seq libraries. All non-synonymous changes were colored in black, synonymous changes in red, and known resistance alleles in green.

### Viral Quasispecies Reveal Both Hard and Soft Selective Sweeps

Single-virion haplotypes should yield deeper insights into how these resistance genotypes evolved. Here, we asked if we could identify whether resistance mutations came from a single source and quickly swept to high frequency (hard sweep), or multiple sources that then grew in frequency independently (soft sweep) [Hermisson and Pennings 2005, Barton 2010, Messer and Petrov 2013]. Note that this question is extremely difficult to address without long-range haplotype information. Hard sweeps are likely to happen with lower mutation rate or extremely strong selection, where adaptive mutations occur one at a time and immediately outcompete other genotypes within the population. Soft sweeps tend to dominate if mutation rates are higher, selection is milder, population is large [Pennings and Hermisson 2006], and multiple lineages carrying advantageous mutations may be present at a time, all increasing in frequency due to the consequent selective advantage [Orr and Betancourt 2001, Innan and Kim 2004]. While HBV is a DNA virus that mutates relatively slowly as compared to RNA viruses, it is also true that Lamivudine exerts a strong selective pressure against viral replication. We also asked if we could identify whether resistance alleles were from de novo mutations or from existing low frequency variants. Adaptation from de novo mutation is usually defined as serial fixation of novel alleles, with just one adaptive allele rising to fixation at a time, whereas adaptation from standing variation often also carries with it multiple pre-existing mutations linked to the advantageous allele [Slatkin and Hudson 1991, Hudson et al., 1994, Barton 1998, Fat and Wu 2000, Durrett and Schweinsberg 2004]. Which model is more relevant is partly determined by population diversity and the presence of pre-existing drug resistant strains. A clonal viral infection, such as a recent or mono-strain infection seeded by very few drug-naïve virions is less likely to carry pre-existing resistance alleles as compared to a mixed infection or a long-term infection that has had time to diversify within the patient. There is also the possibility that a large, highly mutable viral population could theoretically carry all possible mutations in its quasispecies mutant pool at any point in time. Determining the correct model for HBV population evolution will be important for describing and modeling adaptation.

We made use of nearly full genome haplotype information from BAsE-Seq to characterize viral population quasi-species composition before and after drug treatment. Phylogenetic trees built from viral haplotypes revealed three different patterns in how these patients gained viral resistance.

Two patients, P2 and P7, had trees that showed clear mono-clonal gains of resistance, suggestive of hard sweeps (Figure 5,6). Allele frequency changes showed clusters of SNVs that increased in allele frequency together (11 SNVs in P2 spanning the entire 3.2kb sequenced region [Figure 3] and 6 SNVs in P7 spanning 2kb-2.8kb [Figure 4]). Haplotype information confirmed that these were linked SNVs on the same haplotype. 9/11 SNVs in the P2 cluster were within the RT domain of the polymerase gene, and 7/11 SNVs were non-synonymous mutations within a 750bp window. All six SNVs in the P7 cluster were within the RT domain of the polymerase gene, and three were non-synonymous mutations within a 150bp window. These two sweeps with numerous SNVs linked to the resistance allele would support a model of evolution from standing variation. However, these exact combinations of SNVs were not found in the treatment naïve timepoints for either patient. The closest haplotypes found pre-treatment shared just 6/11 SNVs for P2 and 2/6 SNVs for P7. We suggest three possible explanations. First, there could be a detection limit for extremely rare haplotypes in the pre-treatment timepoints. We may simply have failed to sequence them. Second, because our samples came from patient blood samples, latent viral reservoirs outside of the blood stream could be contributing to the viral population. Again, they would be missed by serum samples. Finally, the resistance mutation could have occurred later during the treatment regime by chance, and happened to rise on the background of a viral sequence that already accumulated multiple nucleotide differences from the population consensus. There was in fact a two to three fold difference in nucleotide diversity across the eight patient samples (Table 1), suggesting a range of quasispecies complexity across patients, which may affect observed evolutionary dynamics.

**Figure 5,6.**
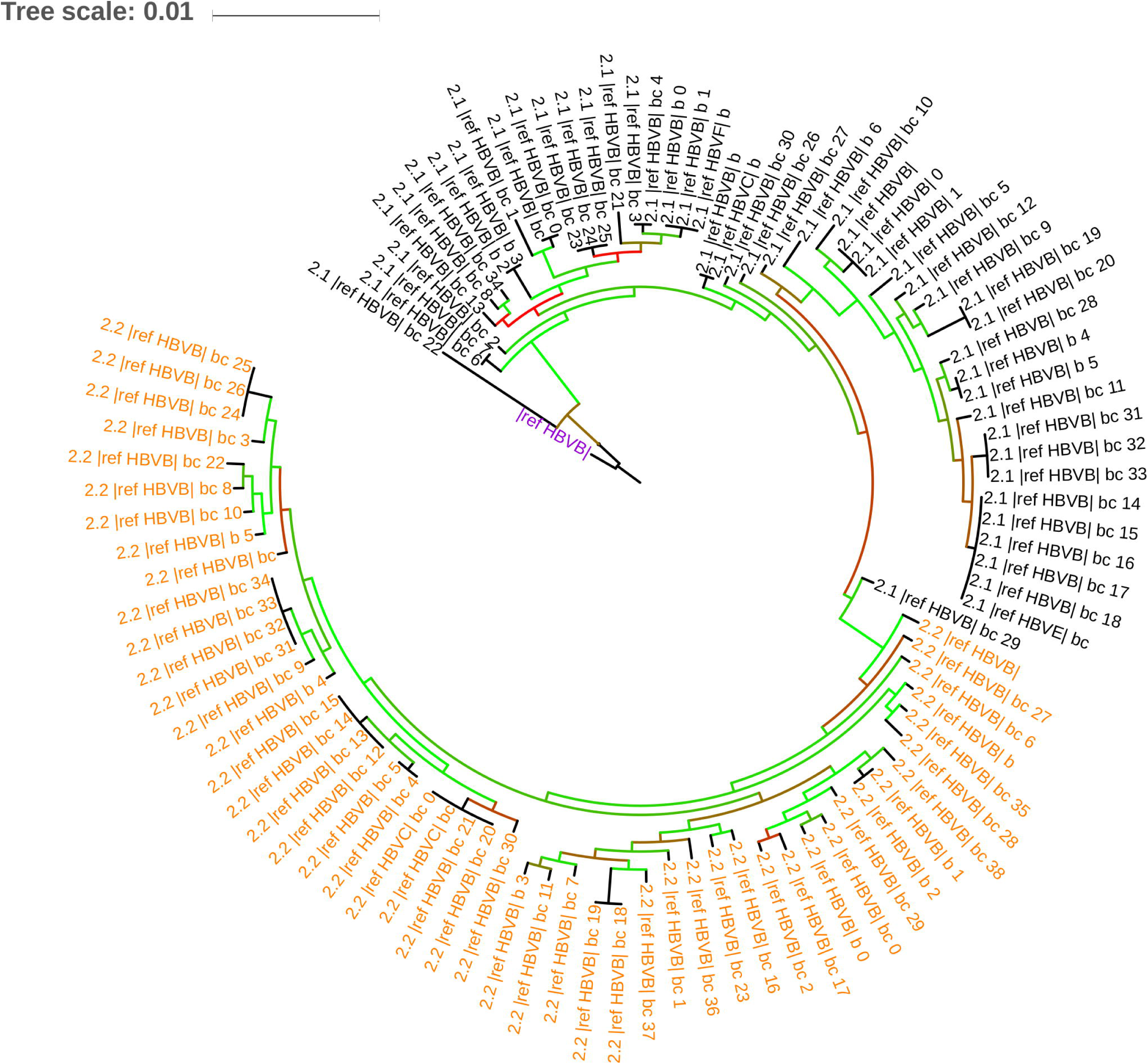
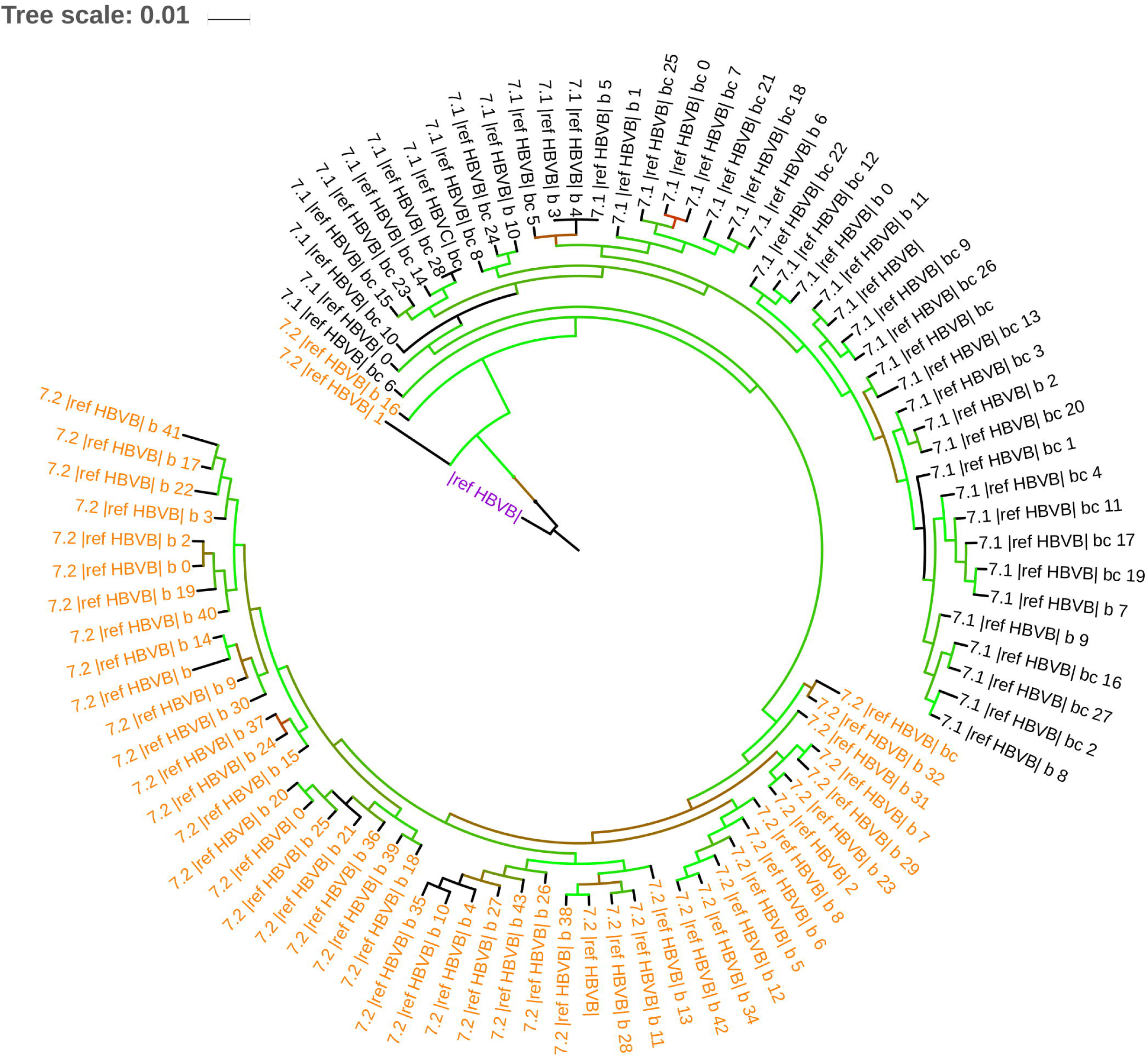
BAsE-Seq approximately maximum-likelihood trees of viral sequences from P2 (Figure 5) and P7 (Figure 6) before (black branches) and after (orange branches) drug resistance. Trees are rooted against reference genome HBV genotype B (purple label). 100 sub-sampled sequences from each timepoint are used, color-coded by their branch labels. Bootstrap values are reflected in branch color, ranging between 0 (red) to 1 (green). Orange indicate sequences post resistance, black indicate sequences pre-resistance. For a full phylogeny of all sequences, refer to (SI Figure 16,17).

The two remaining patients, P1 and P11, had trees showing at least two independent instances of gain of resistance, in other words soft sweeps (Figure 7,8). P1 was highly clonal with just 6 sites shifting in frequency over time (Figure 3), and independently gained M204I and L180M+M204V on two haplotypes. P11 carried the most diverse population out of all four patients, starting as a mixed population of 26% Genotype B and 74% Genotype C (Figure 2). The same resistance allele M204I evolved twice on Genotype C sequences but none on Genotype B sequences, resulting in a resistant viral population that was 100% Genotype C. One lineage further gained the L180M mutation, although it is unclear whether that conferred additional resistance on a M204I background.

**Figure 7,8.**
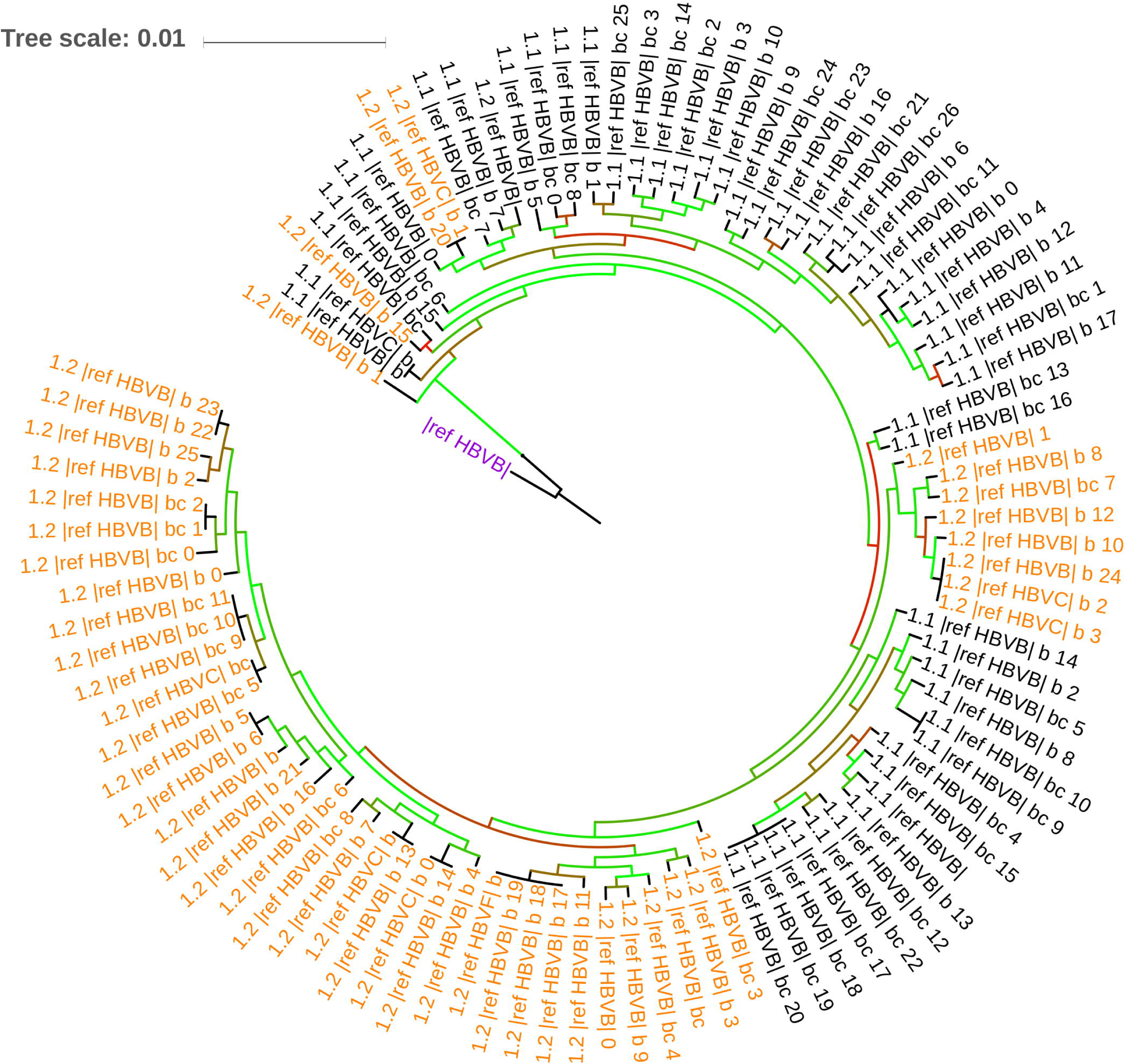
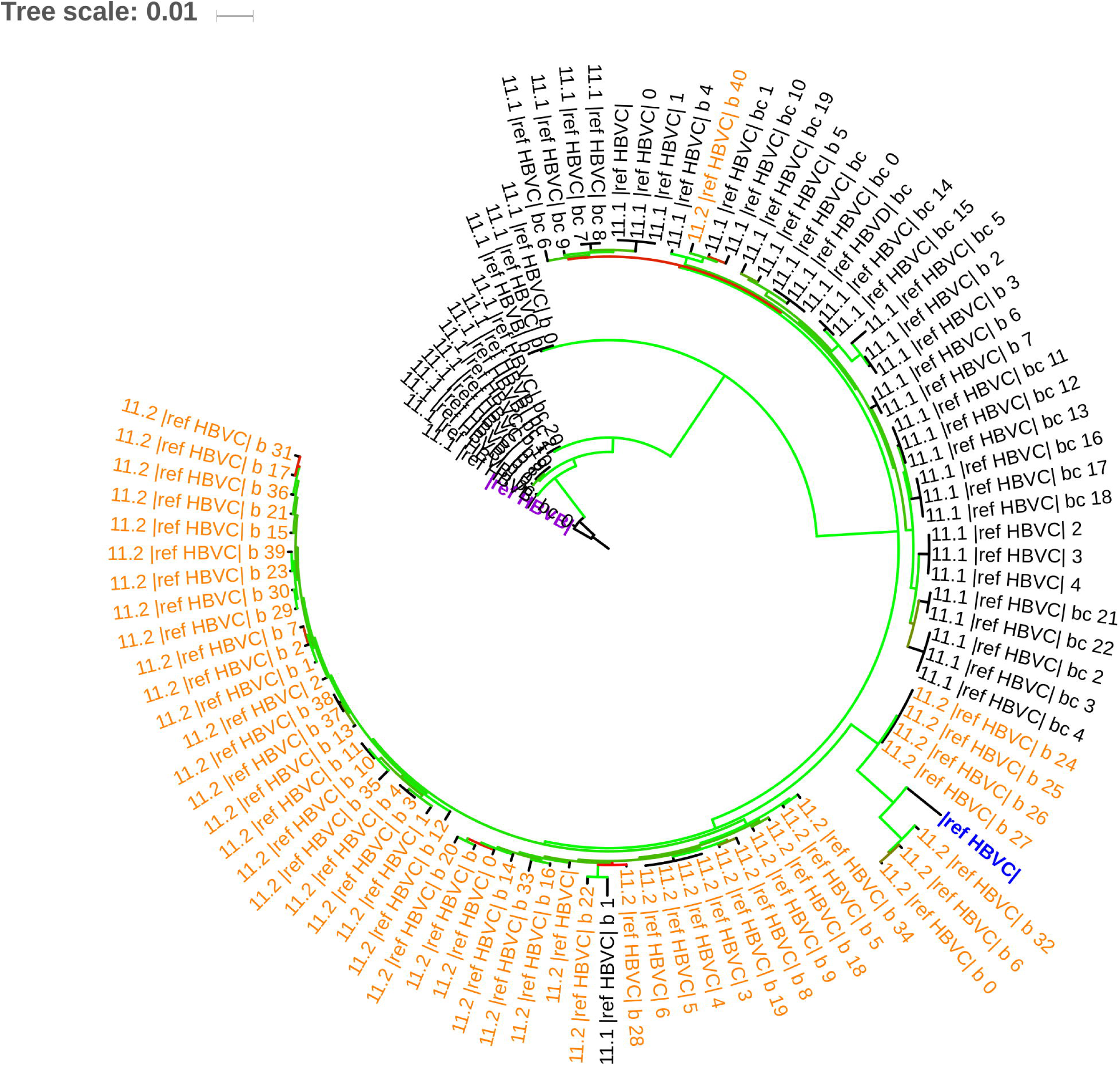
BAsE-Seq approximately maximum-likelihood trees of viral sequences from P1 (Figure 7) and P11 (Figure 8) before (black branches) and after (orange branches) drug resistance. Trees are rooted against reference genome HBV genotype B (purple label). 100 sub-sampled sequences from each timepoint are used, color-coded by their branch labels. Bootstrap values are reflected in branch color, ranging between 0 (red) to 1 (green). Orange indicate sequences post resistance, black indicate sequences pre-resistance. For a full phylogeny of all sequences, refer to (SI Figure 18,19).

### Reconstructing Demographic History

The BEAST analysis for P11 showed effective samples sizes (ESS) of >500, indicating convergence. Sequences came from day 1233 (sample before Lamivudine treatment) and day 0 (sample taken after viral resistance and consequently viral load rebound).

Reconstructed demographic history showed an initial exponential growth phase post infection, followed by a plateau. A sharp dip in effective viral population size (N_e_) then occurred sometime between 750-1350 days prior to day 0. From actual patient viral load information, the population crash post drug treatment occurred sometime around day 1250. There was also a sharp increase in median N_e_ about 150 days prior to day 0 in the skyline plot, which matched actual clinical data almost perfectly (Figure 9). Although the two major changes in population size were well identified, a smaller increase in the viral population size around day 700 was not, possibly indicating some limitations when reconstructing smaller scale changes in population demography, or that these viruses did not contribute to the effective population size. Due to the smaller nucleotide diversity present in the other patient samples (Table 1), runs for P1, P2 and P7 did not coalesce at 1E9 replications (SI Figures 20-22).

**Figure 9.**
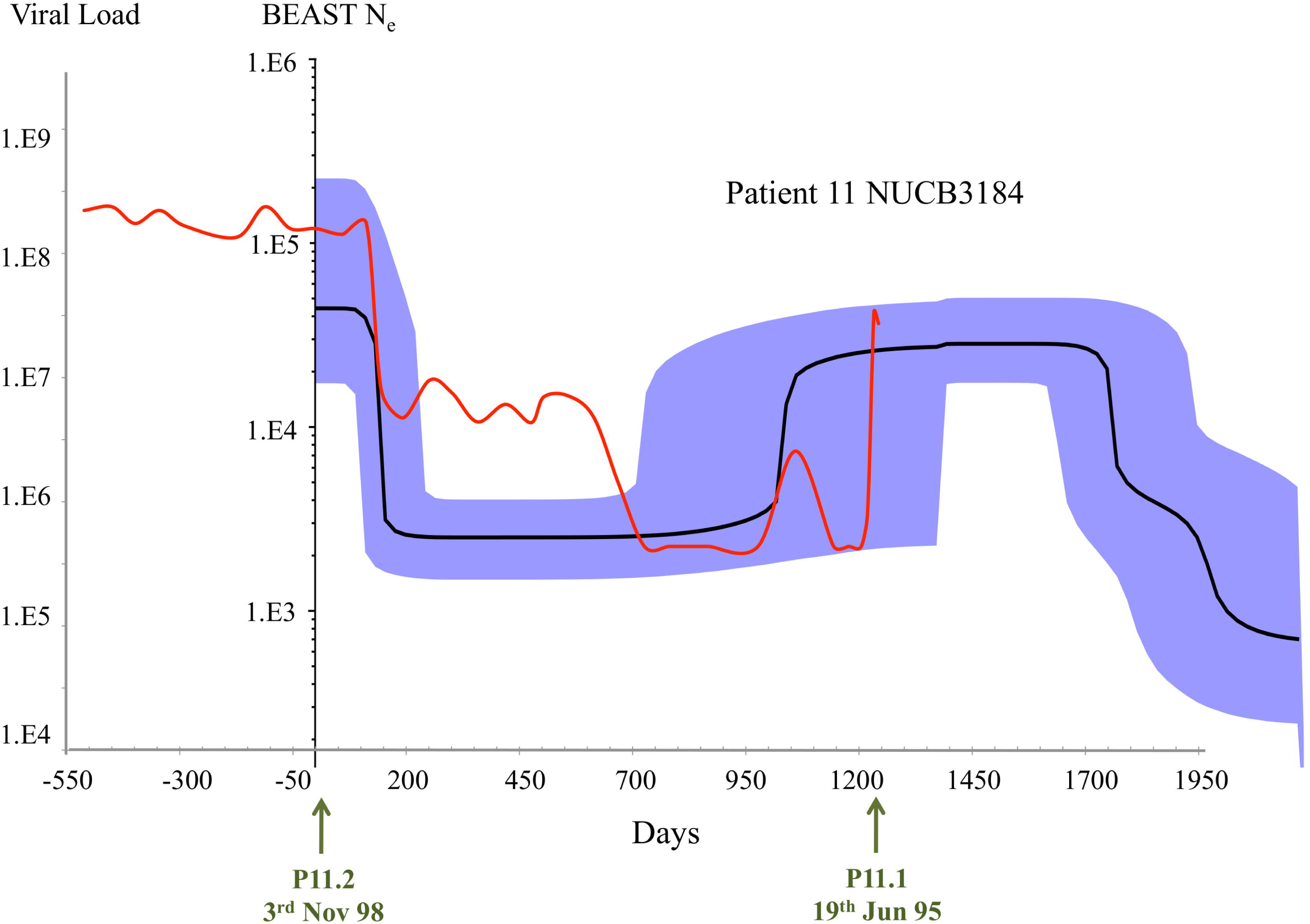
Overlay of BEAST reconstructed demographic history and clinical records of patient viral load (serum) for patient 11. X-axis – Timeline represented in days, going forward in time from right to left. Green arrows point out the two timepoints that were used for single-virion sequencing. Y-axis labeled “Viral load” – corresponding to red line tracing all available records of patient viral load (log10 scale) over time. Y-axis labeled “BEAST N_e_” - Effective population size over time as simulated by BEAST (log10 scale) shown by black line (plotted values are local medians, with 95% highest probability density (HPD) confidence interval colored in blue).

## Conclusions

Haplotype information is vital for revealing hidden population dynamics invisible in standard deep sequencing data. While single-virion sequencing remains technically challenging, we employed two complementary single-virion sequencing platforms to reveal – and cross-validate - such information. We can tell, from nucleotide diversity calculations, the heterogeneity of a population. We can estimate, using hundreds to thousands of single-virion haplotypes, the relative proportions of genotypes and quasispecies present in an infection. We can determine if resistance evolved from a single source, or multiple times independently. Using samples taken at different timepoints, we can begin to explore whether evolution occurs from standing variation or de novo mutations, and how that is linked to quasispecies complexity. While lamivudine resistance is a relatively simple adaptive process with very specific alleles conferring fitness gains, this work shows the potential of applying single-virion sequencing to complex events such as viral response to immune enhancement or viral dynamics during an active HBV flare. It may also be valuable in the study of difficult topics such as cccDNA stability, viral recombination, and viral reservoirs. Single-virion sequencing is therefore a powerful tool for understanding the role of viruses across disease stages of clinical importance.

## Declarations

### Ethics Approval and Consent to Participate

All patients provided written informed consent according to the Declaration of Helsinki, and institutional review board of the participating hospitals approved all study protocols.

### Availability of Data and Material

Raw sequencing reads from each library will be available on NCBI SRA.

### Competing Interests

The authors declare that they have no competing interests.

### Funding

The work presented here was funded by the Agency of Science, Technology and Research (A*STAR) and the grant JCO CDA 13302FG059. YOZ, PA, PFS, SGL, and MH were also funded by the grant titled Eradication of HBV TCR Program: NMRC/TCR/014-NUHS/2015. NN and WFB were also funded by the grant JCO DP 1334k00082.

### Authors’ Contributions

LZH, NN, WFB and MH conceived and designed the experiments. SGL procured the samples. PPKA, PFS, SZH, and LXS constructed the libraries. YOZ, AW, CHL, SH, and MH conducted the analyses. YOZ, PFS, LZH, AW, SH, NN, WFB, and MH drafted and edited the manuscript.

### Authors’ Affiliations

SZH is currently affiliated with Chugai Pharmabody Research Pte Ltd, Singapore 138623. LZH is currently affiliated with Translational Biomarkers, Merck Research Laboratories, MSD, Singapore 138665.

## Acknowledgements

We thank Wendy Soon, Gary Chen and the entire Next Generation Sequencing Platform team at the Genome Institute of Singapore for their support and expertise. We thank Siddarth Signh and the Pacific Biosciences support team for their expertise and invaluable advice and feedback.

